# Mono-allelic epigenetic regulation of bi-directional silencing of RNA Polymerase II polycistronic transcription initiation in *Trypanosoma brucei*

**DOI:** 10.1101/2024.06.21.600114

**Authors:** Rudo Kieft, Laura Cliffe, Haidong Yan, Robert J. Schmitz, Stephen L. Hajduk, Robert Sabatini

## Abstract

Unique for a eukaryote, protein-coding genes in trypanosomes are arranged in polycistronic units (PTUs). This genome arrangement has led to a model where Pol II transcription of PTUs is unregulated and that changes in gene expression are entirely post-transcriptional. Trypanosoma brucei brucei is unable to infect humans because of its susceptibility to an innate immune complex, trypanosome lytic factor (TLF) in the circulation of humans. The initial step in TLF mediated lysis of *T.b.brucei* requires high affinity haptoglobin/hemoglobin receptor (HpHbR) binding. Here we demonstrate that by *in vitro* selection with TLF, resistance is obtained in a stepwise process correlating with loss of HpHbR expression at an allelic level. RNA-seq, Pol II ChIP and run-on analysis indicate HpHbR silencing is at the transcriptional level, where loss of Pol II binding at the promoter region specifically shuts down transcription of the HpHbR containing gene cluster and the adjacent opposing gene cluster. Reversible transcriptional silencing of the divergent PTUs correlates with DNA base J modification of the shared promoter region. Therefore, we show that epigenetic mechanisms, including base J modification, are involved in regulating gene expression via Pol II transcription initiation of gene clusters in a mono-allelic fashion. These findings suggest epigenetic chromatin-based regulation of gene expression is deeply conserved among eukaryotes, including early divergent eukaryotes that rely on polycistronic transcription.

**IMPORTANCE:** The single-cell parasite *Trypanosoma brucei* causes lethal diseases in both humans and livestock. *T. brucei* undergoes multiple developmental changes to adapt in different environments during its digenetic life cycle. With protein-coding genes organized as polycistronic transcription and apparent absence of promoter-mediated regulation of transcription initiation, it is believed that developmental gene regulation in trypanosomes is essentially post-transcriptional. In this study, we found reversible Pol II transcriptional silencing of two adjacent polycistronic gene arrays that correlates with the novel DNA base J modification of the shared promoter region. Our findings support epigenetic regulation of Pol II transcription initiation as a viable mechanism of gene expression control in *T. brucei*. This has implications for our understanding how trypanosomes utilize polycistronic genome organization to regulate gene expression during its life cycle.

## INTRODUCTION

Trypanosomatids are eukaryotic, unicellular parasites, such as *Trypanosoma brucei*, which causes human sleeping sickness. Genomes of these parasites have genes organized into large gene clusters that are transcribed as polycistronic transcription units (PTUs), and messenger RNA maturation occurs by co-transcriptional trans-splicing of spliced leader RNA and polyadenylation (1-4). PTUs can be over 100 kb long and contain genes that are functionally unrelated. Because of this genome arrangement, it is proposed that post-transcriptional regulation is the primary mode of gene expression control (5-7). Control of RNA polymerase II (Pol II) initiation is not an obvious/viable means of individual gene expression control in these early divergent eukaryotes.

An interesting exception to this lack of transcriptional control model is the regulation of Pol I transcription of telomeric PTUs required for the monoallelic expression of the variant surface glycoprotein (VSG), critical for parasite survival in the mammalian host. Unusual for a eukaryote, *T. brucei* contains a multifunctional Pol I able to synthesize a subset of abundant protein coding mRNAs. During infection of the mammalian bloodstream, the exclusive expression of only one VSG gene and the periodic switching of the expressed VSG allows the parasite to evade the host immune system (8,9). While *T. brucei* encodes ∼2500 VSG genes, only one can be transcribed when located within one of ∼15 telomeric VSG expression sites (ESs). These ESs are Pol I transcribed PTUs located adjacent to telomeres on multiple chromosomes and contains a VSG gene and up to 12 expression site-associated genes (ESAGS) (10,11). This highly stringent mono-allelic exclusion ensures that only one telomeric VSG ES is transcribed at a time (12-14). Mechanisms of VSG switching include regulated mono-allelic activation and silencing of these telomeric Pol I transcribed PTUs.

The chromosome core of *T. brucei* is composed of Pol II transcribed PTUs where very little is understood regarding the mechanisms involved in the transcription cycle, including initiation. In fact, sequence-specific core promoters were only very recently identified (15). While regulated Pol II transcription initiation as a mechanism of trypanosome gene expression control has not been demonstrated, several studies have suggested the role of histone post-translational modifications, histone variants, DNA modification and chromatin organization on the regulation of gene expression in trypanosomes (13,16-24). Regarding the regulation of Pol II transcription, trypanosome PTUs are flanked by chromatin marks, including histone post-translational modifications, histone variants, and the DNA modified base J. For example, transcription termination sites at the end of PTUs are enriched in three chromatin marks: the hypermodified DNA base J, and the histone variants H3V and H4V (19,20,25,26). Base J consists of a glucosylated thymidine (27) and has been found in the nuclear DNA of trypanosomatids at almost all Pol II transcription termination sites (26,28). All three epigenetic marks have been shown to play a role in Pol II transcription termination (28-33). Furthermore, for several PTUs in *T. brucei* and *Leishmania*, J/H3V are found to promote Pol II termination prior to the end of the gene cluster, leading to silencing of the downstream genes (29-31). Loss of J/H3V, or knockdown of the base J binding protein 3 (JBP3)-PP1 protein phosphatase complex linking base J with dephosphorylation of Pol II (34-37), results in readthrough transcription and de-repression of the downstream genes. This has suggested regulated ‘premature’ termination of Pol II transcription within a trypanosome PTU as a novel mechanism of regulating the transcription of individual genes.

To address function of premature termination as a viable mechanism of regulatable gene silencing in *T. brucei*, we wanted to explore effects of negative selection for a gene at the end of a PTU on Pol II transcription elongation/termination. A PTU on chromosome 6 contains five genes, with the final gene coding for the haptoglobin/hemoglobin receptor (HpHbR) essential for binding and uptake of the trypanosome lytic factor (TLF)(38-40), an innate cytotoxin associated with a minor subclass of human serum HDLs (41-43). *T. brucei brucei* is unable to infect humans because of its susceptibility to TLF, while other subspecies of trypanosomes that cause human sleeping sickness are capable of circumventing the activity of TLF (44-47). For example, *T. brucei gambiense* has evolved a mechanism that includes down-regulated expression of the HpHbR (48). To study the mechanism of human serum resistance, we treated TLF sensitive *T. b. brucei* Lister 427-221^S^ (expressing the 221 VSG) with progressively higher concentrations of purified human HDL containing TLF (49). A highly TLF resistant population was obtained resulting in two clones, *T. b. brucei* 427-800^R^ and *T. b. brucei* 427-060^R^ (48,49). TLF resistance correlates with complete loss of HpHbR mRNA expression and TLF binding and uptake (48). Although resistance to TLF was a relatively stable characteristic of *T. b. brucei* 427-060^R^, a small fraction of cells spontaneously reverted back to being susceptible to TLF following prolonged culturing in the absence of TLF (48). This 427-060^S^ cell line re-acquired the ability to expresses the HpHbR gene and bind and uptake TLF. Resistance to TLF was confirmed to be caused solely by loss of expression of this receptor by RNAi and by the ability of ectopic expression of the HpHbR gene to restore susceptibility to TLF in the resistant cell lines (48). Therefore, stable and reversible loss of expression of the Hp/Hb receptor, encoded by the HpHbR gene at the end of a Pol II transcribed PTU, can be selected for in *T. brucei*.

The identification of Pol II termination as a viable means for trypanosome gene expression control led us to characterize the mechanism involved in reversible HpHbR gene repression induced by TLF pressure. Here we show that HpHbR silencing is at the transcriptional level, where loss of Pol II binding at the promoter region specifically shuts down transcription of the entire HpHbR containing gene cluster as well the adjacent divergently orientated gene cluster. Reversible transcriptional silencing of the divergent PTUs correlates with DNA base J modification of the shared promoter region. Furthermore, by analyzing additional clonal lines during the TLF selection process we show that resistance to TLF and re-establishing sensitivity is obtained in a stepwise process correlating with repression and de-repression of HpHbR expression at an allelic level. Therefore, reminiscent of the Pol I transcribed telomeric VSG expression sites, epigenetic mechanisms exist to regulate gene expression via Pol II transcription initiation of gene clusters in a mono-allelic fashion.

## RESULTS

### Susceptibility to TLF is dependent on allelic expression of the HbHbR gene

Treatment of *T. brucei* Lister 427-221^S^ with progressively higher concentrations of purified human HDL containing TLF resulted in highly TLF resistant population, represented by the clonal cell line *T. b. brucei* 427-800^R^ (see diagram in Fig 1A)(49). In addition to the resistant phenotype, this cell line expresses a new VSG via duplication and transposition of VSG800 into the 221 ES (49). Cultivation of this resistant cell line in vitro for an extended period led to the isolation of an antigenic variant expressing a new VSG that retained resistance to TLF; *T. b. brucei* 427-060^R^ (48). As we previously described for the 427-800R cell line (49), we confirm that *T. b. brucei* 427-060^R^ is highly resistant to purified TLF and could grow in media containing 25% normal human serum and that this correlates with the complete loss of HbHbR mRNA expression (Fig 1B and C).

**Figure 1.**
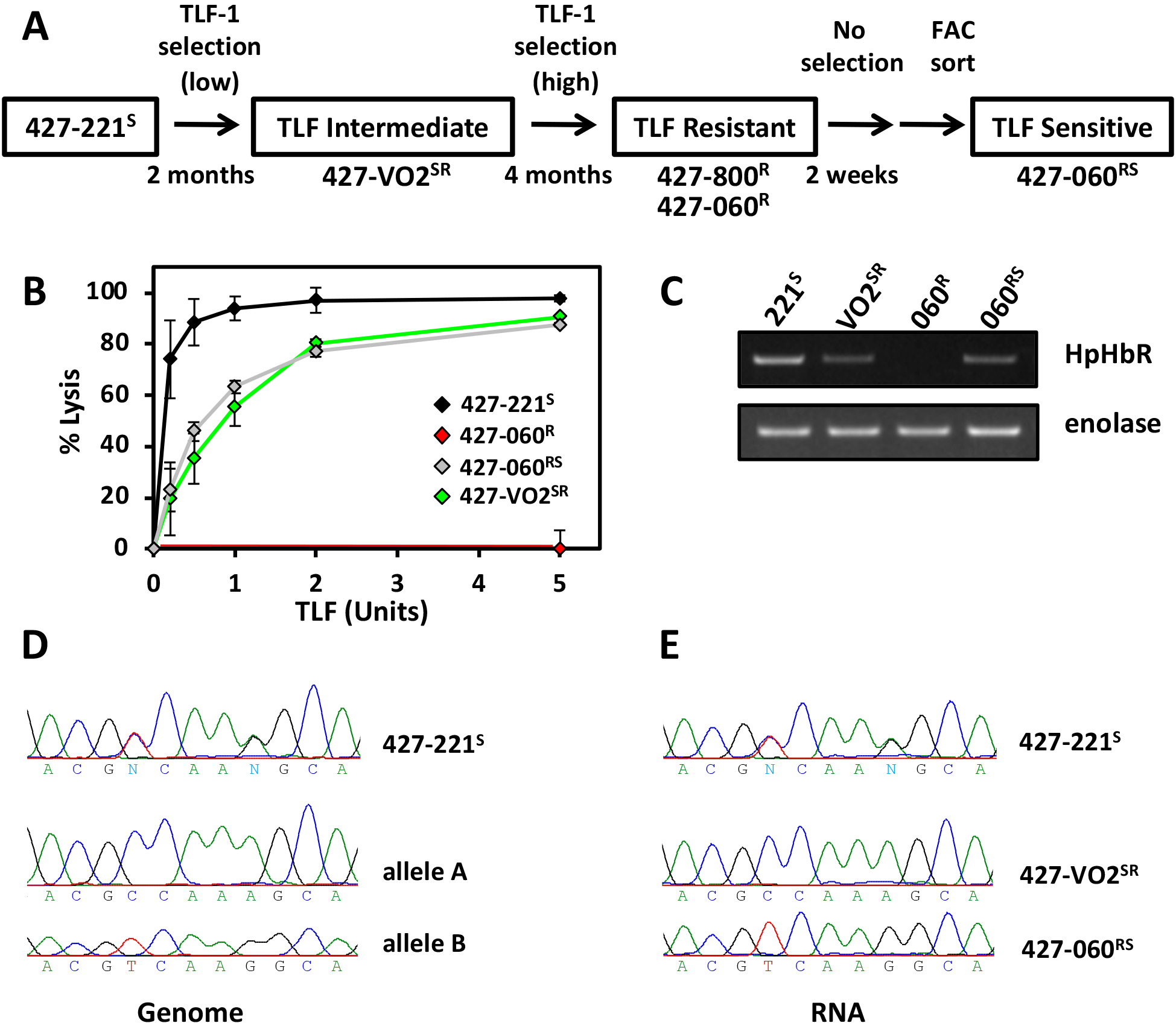
TLF resistance correlates with HpHbR allelic expression. (**A**) Relationship of TLF-1 resistant and susceptible *T. b. brucei* Lister 427 (MiTat 1.2) cell lines used in this study. (**B**) *In vitro* susceptibility of the *T. b. brucei* cell lines to TLF: percentage of cells lysed following incubation with TLF for 2 h at 37° C. *T. b. brucei* 427-221^S^ (black diamond); *T. b. brucei* 427-VO2^SR^ (green diamond); *T. b. brucei* 427-060^R^ (red diamond); *T. b. brucei* 427-060^RS^ (grey diamond). (**C**) RT-PCR analysis of HpHbR (Tb427.6.440) expression for the indicated T. b. brucei 427 cell lines. Enolase expression was used as control. (**D**) Direct sequencing of PCR amplified genomic HpHbR (*T. b. brucei* 427-221^S^) and sub-cloned HpHbR (Tb427.6.440 allele A and allele B) demonstrates the presence of 2 distinct allelic copies of HpHbR. PCR amplicon allows analysis of the C/T SNP and the A/G SNP at position 970 and 1106, respectively. (**E**) Direct sequencing of RT-PCR amplified HpHbR mRNA from *T. b. brucei* 427-221^S^ (expression of allelic copies A and B), *T. b. brucei* 427-VO2^SR^ (expression of allelic copy A) and *T. b. brucei* 427-060^RS^ (expression of allelic copy B).

Although TLF resistance was a relatively stable characteristic of *T. b. brucei* 427-060^R^, a small fraction cells were observed to spontaneously revert back to being susceptible after *in vitro* growth without TLF pressure for a few weeks and isolated following FACs enrichment based on TLF uptake (48). To reflect the intermediate TLF resistance-sensitive phenotype (see below) and being derived from a highly resistant cell line this 427-060^S^ clonal line is now referred to as *T. b. brucei* 427-060^RS^. Upon closer examination of these cells using a more sensitive TLF lysis assay, we observe that *T. b. brucei* 427-060^RS^, while high susceptible to TLF killing, still maintains some level of TLF resistance when compared to the parental *T. b. brucei* 427-221^S^ (Figure 1B). This intermediate TLF resistance phenotype correlates with intermediate level of HpHbR gene re-expression (Figure 1C). Similarly, we have isolated a clonal line, *T. b. brucei* 427-VO2^SR^, from an earlier point along the in vitro TLF selection scheme (Figure 1A) that has similar intermediate levels of TLF resistance (Figure 1B) and corresponding intermediate level of HpHbR gene expression (Figure 1C).

The apparent ∼50% level of HpHbR gene expression in the two clones exhibiting intermediate TLF related phenotypes, *T. b. brucei* 427-VO2^SR^ and 427-060^RS^, could be explained by allelic control of gene expression. An interesting idea that could be addressed once we noticed 8 SNPs within the HpHbR gene in the *T. brucei* 427 genome that would allow us to follow specific allelic expression (Table S1A). Sequencing of PCR clones from genomic DNA confirmed the SNPs in the HpHbR gene allowing us to distinguish between the two alleles we refer to as allele A and B (Figure 1D and Table S1A). Sequencing of RT-PCR amplified HpHbR RNA from the two intermediate phenotypic clones indicates distinct alleles expressed in each; allele A in *T. b. brucei* 427-VO2^SR^ and allele B in *T. b. brucei* 427-060^RS^ (Figure 1E and Table S1A). Both allelic copies are equally independently functional, as we demonstrate ectopic expression of each allelic copy of the HpHbR gene rescues TLF binding (reflected by Hp binding) and TLF lysis in the TLF resistant cell line 427-800^R^ (Figure 2A-C). Interestingly, the increased expression of Allele B from the tubulin array compared to the native locus in wild type 427-221^S^ cells, resulted in an increased level of susceptibility to TLF lysis (Figure 2C and D). Both allelic copies are also maintained throughout the TLF selection procedure, confirming that the selective expression of distinct alleles in the two intermediate phenotypic clones represents monoallelic expression rather than reflecting loss of heterozygosity or mutagenesis at the HpHbR gene locus. Following the allele that is expressed in each intermediate phenotype clone by mRNA RT-PCR sequencing of the entire HpHbR ORF (compared with the parental with both alleles expressed) indicates that allele B is initially silenced (VO2^SR^), but after passing thru the silencing state where both alleles are silenced (060^R^), allele B is the one activated in the subsequent intermediate state (060^RS^) (Figure 1E and Table S1A). Thus, the previously silent allele B is maintained as it is the allele that is selectively activated in the final TLF sensitive intermediate (060^RS^). Taken together, we have shown that the degree of TLF resistance (and sensitivity) correlates with degree of HpHbR gene expression, and that intermediate levels of gene expression is linked with allelic regulation.

**Figure 2.**
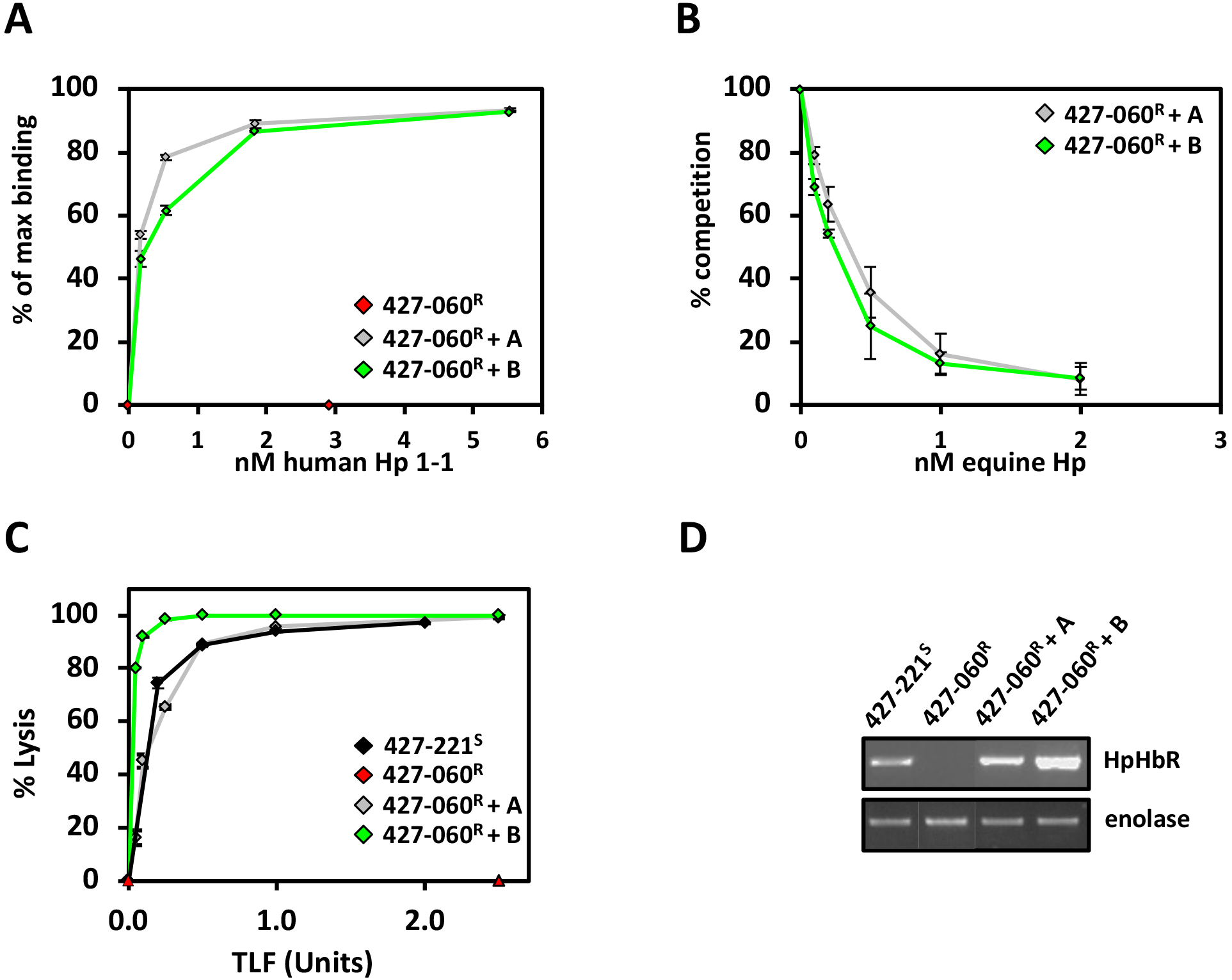
Allelic copies of HpHbR are functional in haptoglobin-hemoglobin binding and TLF lysis. (**A**) Saturation binding at 3°C with increasing concentrations of Alexa-488 labeled human haptoglobin (Hp 1-1) in the *T. b. brucei* 427-060^R^ cells ectopically expressing Tb427.6.440 allelic copy A or Tb427.6.440 allelic copy B from the tubulin array. (**B**) Competition for Alexa-488 human HpHb binding with unlabeled equine HpHb in the 427-060^R^ transfected cell lines as in A. (**C**) *In vitro* TLF assay. *T. b. brucei* 427-221^S^ is shown as a black triangle; *T. b. brucei* 427-060^R^ a red triangle; 427-060^R^+ HpHbR Allele A rescue a grey triangle; and 427-060^R^+ HpHbR Allele B rescue a green triangle. (**D**) Levels of HpHbR gene expression in the 427-060^R^ transfected cell lines as in A were determined by RT-PCR and compared with untransfected 427-060^R^ and levels from the actively transcribed native locus in 427-221^S^ cells. Enolase provides a positive control. The differing levels of allele A and B HpHbR expression from the construct inserted within the tubulin array, presumably represent clonal variations commonly seen in *T. brucei* genome integrative constructs.

### Downregulation of HpHb mRNA expression is due to silencing Pol II transcription initiation

Allelic control of HpHbR gene expression suggests regulation at level of Pol II transcription. For example, Pol II termination now occurring prior the HpHbR gene in the PTU. Consistent with this, the lack of HpHb mRNA in the 427-060^R^ resistant cell line is presumably not due to any changes in RNA stability as we show that ectopic expression of the HpHbR (with native UTR sequences) in 427-060^R^ cells has similar mRNA half-life as endogenous HpHbR mRNA in the parental sensitive cell line (427-221^S^) (Figure S1). To characterize localized changes in the transcriptome at the HpHbR gene locus, and address premature termination within the PTU, we performed stranded mRNA-seq to compare the expression profiles of TLF sensitive and resistant cell lines (Figure 3A and B). Differential expression analysis (DESeq2 module) revealed changes in mRNA expression of ∼300 genes distributed throughout the *T. brucei* genome using a cut-off value of a 3-fold log2 in the 427-060^R^ compared to the sensitive parental 427-221^S^ cell line (Figure S2). Changes in expression such as this can be expected among cell lines grown for such an extended period of time (∼6 months). This includes changes that reflect the VSG recombinational switch that occurred during the generation of the 427-060^R^ cells (48,49). Strikingly, however, we see a clustering of significant gene downregulation at the HpHbR gene locus (Figure 3A and B). Among the only seven genes with altered expression using a cut-off value of a 6-fold log2, six are downregulated genes in 427-060^R^ cells representing all five genes in the HpHbR gene array (including the HpHbR gene itself) and one of the five genes in the adjacent opposing PTU (Figure 3B, top). While two of the other genes in the adjacent PTU are also downregulated, albeit only 2-3-fold log2, another two genes appear not be affected. However, the unaffected genes are multicopy genes that could alter the RNA-seq read mapping. Consistent with allelic specific de-repression, the silencing of these genes in the TLF resistant cell line (427-060^R^) appears to be partially restored in the revertant, partially sensitive 427-060^RS^ cell line (Figure 3B, bottom). Specific changes in the RNA expression of representative genes in the opposing PTUs, and no change in expression for a gene in another PTU further downstream, are confirmed by RT-PCR (Figure 3C). To examine whether the reversible silencing of the HpHbR was in fact due to decreased transcription of the entire PTU, as opposed to post-transcriptional mechanisms, we performed nuclear run-ons. To study closely related isogenic cell lines, as well as avoid cells that have undergone a VSG switching event, we compared TLF resistant 427-060^R^ to the sensitive revertant 427-060^RS^. Consistent with decreased Pol II transcription of the divergent PTUs, the signal from elongating Pol II is specifically lost from the entire region representing the HpHbR PTU and the opposing adjacent PTU in the TLF resistant 427-60^R^ cells (Figure 4A and B). The lack of Pol II transcription of this bi-directional PTU region is in contrast to other regions of the genome, including a unique PTU further downstream (represented by probes 15-17)(Figure 4A and B). Transcription of this region is then re-established in the sensitive 427-060^RS^, albeit at significantly lower levels than genes in the PTU further downstream (Figure 4B). This apparent locus specific lower Pol II activity in the 427-060^RS^ cells may reflect mono-allelic transcription of the divergent gene arrays. SNPs were identified for other genes in the 427 genome including the 6.360 gene located in the adjacent opposing PTU that appears to be co-regulated with the HpHbR gene array and the 6.650 and 6.740 genes located further downstream within an unrelated gene array (Table S1B-D). As we showed for HpHbR gene expression in the intermediate phenotypic cell lines (427-VO2^SR^ and 427-060^RS^), RT-PCR sequencing of mRNA indicates distinct alleles are expressed for the 6.360 gene located in the adjacent opposing PTU (Table 1SB) consistent with decreased levels of mRNA (Figure 3B). The alternating selective expression of distinct alleles for genes (6.440 and 6.360) in each adjacent divergent PTU during the TLF selection procedure further confirms monoallelic exclusion of the entire co-regulated divergent PTU locus. As shown for the HpHbR gene, the silent 6.360 gene allele in the VO2^SR^ cell line is maintained as it is the allele that is selectively activated in the final TLF sensitive intermediate (060^RS^) after passing thru the silencing state where both alleles are silenced (060^R^) (Table S1B). In contrast, consistent with the lack of change in mRNA expression levels, both alleles of the 6.650 and 6.740 gene in the downstream unrelated gene array are continued to be expressed (Table 1SC and D). Taken together, the results suggest that TLF pressure results in allelic silencing Pol II transcription initiation of the adjacent PTUs, rather than specific regulation of the HpHbR gene via premature termination.

**Figure 3.**
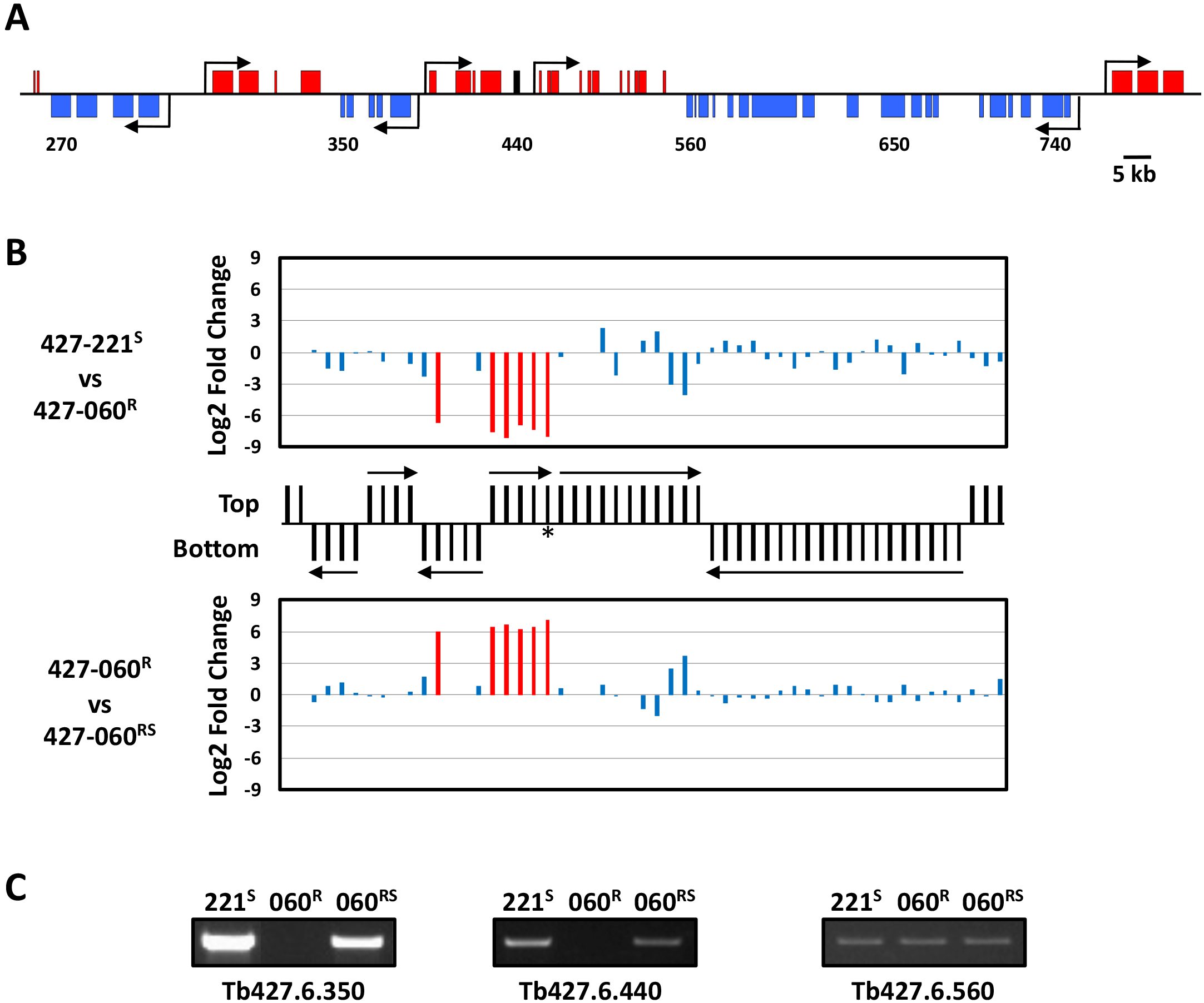
Silencing of HpHbR expression in TLF resistant *T. brucei* is associated with regulated expression of the entire PTU. (**A and B**) Changes in transcript levels based on RNA-seq reads during in vitro TLF selection and generation of *T. b. brucei* 427-060^R^ and 427-060^RS^ cell lines. (**A**) Schematic representing a core section of chromosome 6 corresponding to the divergent HpHbR transcription unit and the adjacent convergently arranged Pol II transcribed units. The HpHbR gene is highlighted in black. Arrows represent transcription start sites and the direction of transcription. Gene numbers are shorthand for genome annotation. For example, 440 refers to Tb427.6.440 (HpHbR). (**B**) Gene map of the core section of chromosome 6 is shown in the center of the figure that includes the region shown in A. mRNA coding genes on the top strand are indicated by black lines in the top half of the panel, bottom strand by a line in the bottom half. Genes on the top strand are transcribed from left to right and those on the bottom strand are transcribed from right to left. Arrows indicate the direction and limits of PTU transcription. Asterisk indicates the HpHbR gene. Above, changes that occur during conversion of *T. b. brucei* 427-221^S^ to *the* resistant 427-060^R^. Genes that are upregulated or downregulated >3-fold Log2 are highlighted in red. Below, changes that occur during conversion of TLF resistant *T. b. brucei* 427-060^R^ to sensitive 427-060^RS^. (**C**) RT-PCR analysis of expressed Tb427.6.350, Tb427.6.440 (HpHbR) and Tb427.6.560 from the indicated *T. b. brucei* 427 cell lines.

**Figure 4.**
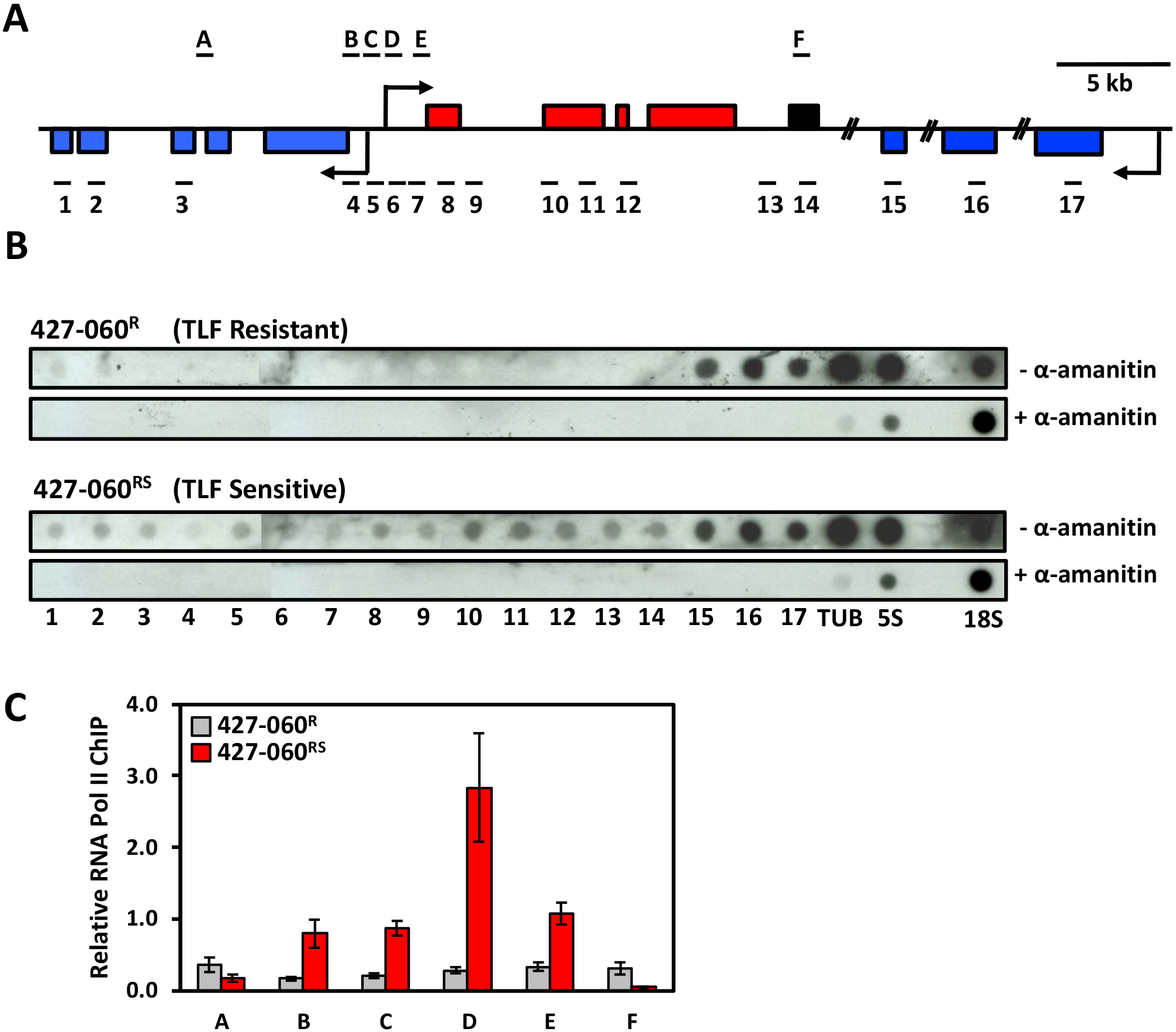
Silencing of HpHbR via PTU transcriptional regulation of the bi-directional Pol II promoter. (**A**) A schematic of the divergent HpHbR transcription unit and the adjacent convergently arranged unit as in Figure 3A. The HpHbR gene is highlighted in black. The locations of the DNA probes (1-17) used in the nuclear run-on analysis are indicated underneath (see Table S2 for oligonucleotides used). The location of the DNA fragments (A-F) used in qPCR analyses is shown above (see Table S2 for oligonucleotides used). Arrows indicate the putative RNA Polymerase II transcription start sites. (**B**) Nuclear run-on analysis of *T. b. brucei* 427-060^R^ and *T. b. brucei* 427-060^RS^. DNA probes were spotted onto nitrocellulose filters using a Dot-Blot apparatus. Tub, β-tubulin; 5S, 5S RNA; 18S, 18S RNA; represent controls for RNA Polymerase II, III and I transcription, respectively. Nascent transcripts were radio-labelled in the absence and presence of α-amanitin (see Materials and Methods). (**C**) Decreased transcription of the divergent PTU correlates with decreased Pol II occupancy at the bi-directional promoter region. Decreased RNA polymerase II occupancy at the Pol II bi-directional promoter in TLF resistant with decreased PTU transcription. Pol II occupancy within the divergent SSR of *T. b. brucei* 427-060^RS^ (grey bars) and *T. b. brucei* 427-060^R^ (red bars) cells determined by anti-Pol II ChIP/qPCR. The schematic in A represents the divergent SSR, where horizontal lines (A-F) indicate the approximate positions of amplicons. All the data are normalized to the SL-RNA gene, and no antibody control was used as a negative control. Data represents three independent experiments. Standard deviations are indicated with error bars.

Eukaryotic gene expression begins with recruitment of the transcription machinery to a gene promoter and formation of a preinitiation complex composed of Pol II and general transcription factors. Analysis of Pol II ChIP datasets and individual core promoter sequence activity indicates the 5-10 kb regions between the 5’ ends of two divergently orientated PTUs in *T. brucei*, so-called divergent strand-switch region (SSR), contain two oppositely oriented core promoters that initiate transcription in a directional manner (15,22). To evaluate the impact of negative TLF pressure on formation of the preinitiation complex at the promoter region for the divergent PTUs, we investigated the changes in Pol II promoter occupancy upon transcriptional de-repression of the HpHbR PTU by ChIP using specific antibodies to the large subunit of *T. brucei* Pol II. Analysis of Pol II occupancy within the divergent SSR of TLF resistant 427-060^R^ cells using ChIP/qPCR indicated the lack of Pol II in the entire region (Figure 4C), consistent with the lack of Pol II transcription of the divergent PTUs. Upon de-repression of transcription in 427-060^RS^ cells we find Pol II binding within the divergent SSR with highest levels of Pol II at the previously mapped Pol II enriched sequence in WT *T. brucei* cells (Figure 4C and Figure S3). Therefore, development of TLF resistance in 427-060^R^ cells led to an apparent loss in Pol II occupancy at these sites within the divergent SSR and corresponding loss of nascent mRNA formation within the adjacent polycistronic units. These results suggest that the loss of HpHbR expression during the selection of TLF resistance is due to transcriptional repression of a promoter via regulated formation of the Pol II preinitiation complex that can be reversed to reactivate allelic transcription of the divergent PTUs.

### Epigenetic regulation of bi-directional Pol II transcription initiation

To characterize changes in the chromatin environment of the Pol II promoter resulting from TLF selection in *T. brucei*, and explain the transcriptional silencing of the adjacent PTUs, we performed FAIRE and H3 ChIP analysis of the divergent SSR (Figure 5A and B). FAIRE analysis identifies nucleosome-depleted or naked regions and has been utilized to characterize the chromatin landscape of Pol I, Pol II and III loci in trypanosomes (16,50,51). When formaldehyde-cross-linked and sheared chromatin is extracted, fragments that were naked or contained loosely associated histones or other proteins are enriched in the aqueous phase. We show no significant change in abundance of the soluble fraction of chromatin or levels of Histone H3 at the promoter region of 427-060^R^ and 427-060^RS^ cells (Figure 5 A and B). Suggesting no significant global changes in chromatin structure (nucleosome abundance) in the 427-060^R^ cells to explain the silencing of Pol II initiation.

**Figure 5.**
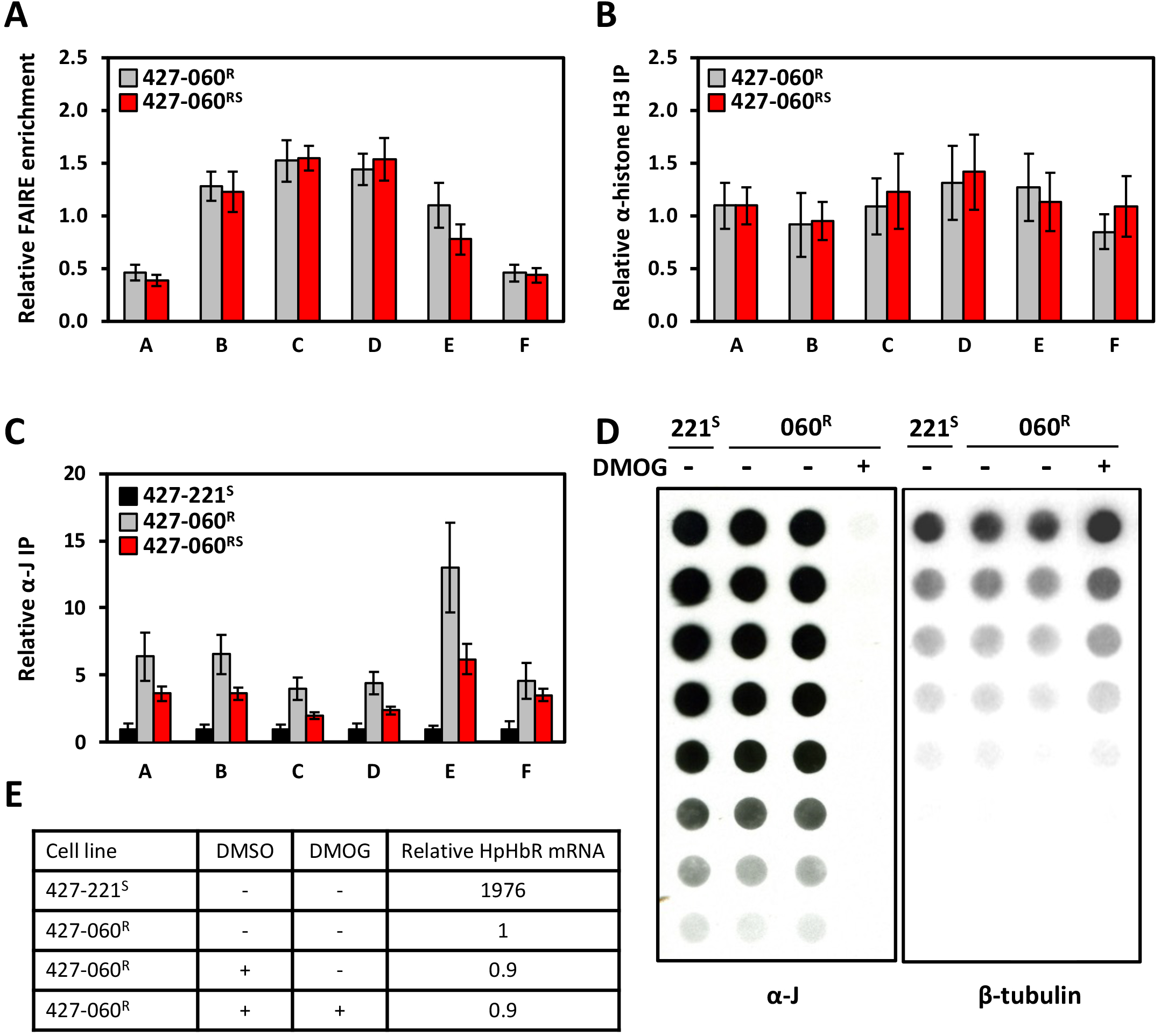
Ablation of RNA Polymerase II binding and increased levels of base J correlates with the loss of bi-directional transcription activity. (A) FAIRE analysis of the divergent Strand Switch Region of the HpHbR transcription unit in *T. b. brucei* 427-060^RS^ (grey bars) and *T. b. brucei* 427-060^R^ (black bars). FAIRE enrichment is shown as the difference in aqueous phase DNA between cross-linked and non-cross-linked samples. The letters refer to PCR regions that correspond to the divergent SSR analyzed as shown in Fig 4A. All calculated values are normalized versus Tb427.6.740 gene sequence control. Results show the average from three independent experiments, with the standard deviations indicated with error bars. (**B**) Histone H3 distribution. Chromatin Immune Precipitation (ChIP) in *T. b. brucei* 427-060^RS^ (grey bars) and *T. b. brucei* 427-060^S^ (black bars), using an antibody against Histone H3, analyzed by qPCR. Relative IP values are calculated versus Tb427.6.740. qPCR fragments as in 4A. (**C**) Base J levels determined by anti-J ChIP. Immune precipitation in *T. b. brucei* 427-221^S^ (black bars), *T. b. brucei* 427-060^R^ (red bars) and *T. b. brucei* 427-060^RS^ (grey bars), using an antibody against the modified DNA base J, analyzed by qPCR. Relative IP values are calculated versus Tb427.6.740. qPCR fragments as in 4A. (**D**) Inhibition of base J synthesis in the TLF resistant cell line. Incubation of *T. b. brucei* 427-221^S^ (221^S^) and *T. b. brucei* 427-060^R^ (060^R^) in the absence and presence of 1 mM DMOG results in global loss of base J from the genome. (**E**) HpHbR expression following the loss of base J in the TLF resistant cell line. qRT-PCR analysis of HpHbR expression in *T. b. brucei* 427-221^S^ and *T. b. brucei* 427-060^R^ cells following the loss of base J by incubation with DMOG. The mRNA expression level in the 427-060^R^ is set to one.

To determine the distribution of base J at this divergent Pol II promoter region and changes associated with reversible silencing of the HpHbR locus, we performed anti-J ChIP-PCR. Consistent with our previous analysis by anti-J ChIP-seq (26), very little base J is present in this SSR of WT *T. brucei* (427-221^S^) cells (Figure 5C). Upon selection of TLF resistance (427-060^R^) we see a significant increase of base J in the promoter region for the HpHbR PTU (Figure 5C) that correlates with the low level of Pol II occupancy (Figure 4C) and transcriptional silencing of the divergent PTUs (Figures 3B and 4B). Conversion of these cells back to TLF sensitive cells (427-060^RS^), led to a decrease in base J that correlates with increased Pol II occupancy and transcriptional activation of the divergent PTUs. Interestingly, base J levels at the promoter region in the sensitive 427-060^RS^ cells do not fully decrease to initial levels present in WT parental cells (427-221^S^), potentially reflecting mono-allelic repression that remains in the 427-060^RS^ cells. Inhibition of thymine hydroxylase activity of JBP1 and JBP2 in the J biosynthesis pathway with DMOG leads to the loss of base J from the genome during subsequent DNA replication (52). Growth of the 427-221^R^ cells in DMOG led to complete loss of base J, but with no effect on HpHbR gene expression (Figure 5D and E). These results suggest that while base J correlates with silencing Pol II initiation at the HpHbR PTU promoter region, its presence is not needed for maintenance of transcriptional silencing.

## DISCUSSION

Organization of protein-coding genes in large PTUs provides no obvious way of regulating the transcription of individual genes and led to the hypothesis that trypanosomes lack selective transcription initiation, with all PTUs being transcribed at similar rates. Accordingly, Pol II transcription has largely been considered to be constitutive and initiation largely dispersed from undefined sites within regions, such as the SSRs, located at the 5’ end of PTUs. However, sequence specific promoters that drive accurate initiation of transcription in an orientation-dependent manner were recently identified in *T. brucei* (15). These studies further suggested, along with earlier studies (22), that coordination between promoter sequence and chromatin context may allow mRNA levels to be regulated via Pol II initiation in trypanosomes. This is supported by our previous work in *T. cruzi* demonstrating suppression of Pol II transcription initiation by base J regulation of chromatin structure (16,17). Furthermore, epigenetic control of ‘premature’ Pol II termination within PTUs of *T. brucei* and *L. major* demonstrated a mechanism for control of individual gene expression within a polycistronic system (29-31,37). We now show that growth of *T. brucei* under negative selection pressure of the human TLF toxin reduces the expression of functional TLF receptor protein (HpHbR) via epigenetic regulation of Pol II binding at the promoter region, silencing the transcription of two PTUs representing 10 genes. Not only do these studies confirm that sensitivity (and resistance) of 427 *T. b. brucei* to the human TLF toxin is due to a single HpHbR gene present at the end of a Pol II transcribed gene array, but also support epigenetic regulation of Pol II PTU initiation as a viable mechanism of gene expression control in *T. brucei*.

We have previously demonstrated that Pol II transcription initiation at divergent SSRs in *T. cruzi* is regulated by base J (16,17). The loss of base J from these regions leads to decreased nucleosome abundance, increased Pol II occupancy and increased Pol II transcription of the adjacent polycistronic units (16,17). Therefore, we proposed that like the mechanism of epigenetic modifications in other eukaryotes, the presence of base J at promoters alters chromatin structure and changes the accessibility of DNA binding proteins, including RNA Pol II, thereby regulating transcription initiation and trypanosome gene expression. However, similar studies of base J function in *T. brucei* failed to illustrate a role in Pol II promoter function. Loss of base J in *T. brucei* led to changes in Pol II termination at the 3’ end of PTUs with no apparent effects on Pol II initiation (29,30). Studies such as these have suggested that the synthesis of base J in BS form *T. brucei* is to help restrict Pol II transcription to within the chromosome core and help control the expression of genes critical during infection of the mammalian bloodstream. We have recently demonstrated that a consequence of Pol II termination defects in *T. brucei* is pervasive Pol II transcription and the expression of the silent Pol I transcribed telomeric VSG ESs (37). Suggesting that a key function of base J in *T. brucei* is tight control of Pol II transcription termination at the PTUs in the chromosome core and help maintain mono-allelic VSG exclusion during infection of the mammalian bloodstream. This idea may help explain why base J synthesis is restricted to the BS life-stage of *T. brucei* and ∼50% of base J is localized to telomeric regions, including DNA repeats immediately upstream of the Pol I VSG ES promoters.

We have also described the role of base J and H3V in regulation of ‘premature’ Pol II termination within PTUs of *T. brucei* and *L. major* as a mechanism for control of individual gene expression within a polycistronic system (29-31,34,36). Therefore, we were surprised that the independent clones analyzed here silenced HpHbR expression following TLF pressure by silencing Pol II transcription initiation. We envisioned that regulated Pol II termination prior to the HpHbR gene would be favored since it allows selective silencing of the final gene of the array. In contrast, silencing Pol II initiation would affect all genes in the PTU. Furthermore, it is difficult to imagine a mechanism to independently regulate promoter systems at divergent PTUs in these genomes. In fact, we find here that silencing of HpHbR gene array is linked to the silencing of the opposing PTU, and as a consequence, downregulating all genes within both arrays. The HpHbR locus is unique in that both PTUs are very small with 5-6 genes that are not essential for *in vitro* growth of bloodstream form *T. brucei* in semidefined media. Trypanosome PTUs usually contain dozens to hundreds of genes. Selection against expression of a gene at the end of a larger PTU may lead to different results. Monoallelic transcription of both PTUs is reflected by SNPs in cell lines that display intermediate levels of HpHbR mRNA expression and corresponding degrees of TLF binding and lysis, compared with the parental TLF sensitive 427-221^S^ cells. At the chromatin level this is reflected by intermediate levels of base J at the Pol II promoter. Therefore, reminiscent of the Pol I transcribed telomeric VSG expression sites; epigenetic mechanisms exist to regulate gene expression via Pol II transcription initiation of gene clusters in a monoallelic fashion in *T. brucei*. Strict monoallelic control of the telomeric VSG PTUs makes sense for antigenic variation and the ability of *T. brucei* to effectively evade the host immune system. An essential role for monoallelic control of the chromosome core is less obvious. In the case of the HpHbR gene silencing in response to TLF pressure we describe here, monoallelic control is not driven by any functional differences between the gene alleles. While there are 8 SNPs in the HpHbR ORF there is no apparent functional difference between the allelic copies. We show both are equally able to function as TLF receptor in vivo. Rather, we assume this represents an example of random monoallelic expression, generated through a stochastic process. The initial random choice between the two alleles is followed by a stable transmission of monoallelic expression. Likely providing a convenient and rapid way to decease-increase transcript levels, analogous to X-chromosome inactivation in mammals, in response to TLF pressure. It remains to be seen if genes with allelic non-synonymous SNPs exist where monoallelic control, and a mixed population of cells in which different alleles are active, provides a benefit to the *T. brucei* lifecycle. It will be interesting to examine whether these epigenetic mechanisms regulating Pol II initiation and termination in a polycistronic array are ever utilized in vivo during the parasite life cycle.

## MATERIALS AND METHODS

### *In vitro* growth, Generation and Transfection of *T. b. brucei* Cell Lines

Bloodstream form *T. b. brucei* Lister 427 (MiTat 1.2) were used in these studies. TLF resistant *T. b. brucei* 427-800^R^ and 427-060^R^ and sensitive 427-060^RS^ cells were generated as previously described (49). During the in vitro generation of the TLF resistant 427-800^R^ line, a population of cells resistant to intermediate levels of TLF were obtained (427-12) and were cloned by limiting dilution. This clone is designated *T. b. brucei* 427-VO2^SR^, based on its intermediate TLF sensitivity, being derived from a highly sensitive cell line and expressing the VO2 VSG. Transfections were performed using the Amaxa electroporation system (Human T Cell Nucleofactor Kit, program X-001).

### Ectopic Expression of the HpHb Receptor

The Hp/Hb receptor ORF was PCR amplified from *T. brucei* 427-221^S^ and cloned into the pTub-phleo construct as previously described (48). Sequencing confirmed constructs representing allele A and allele B with Actin and Tubulin UTR sequences were generated. The Hp/Hb receptor alleles were transfected in *T. brucei* 427-060^R^ cells for analysis of TLF binding and TLF lysis. Another set of pTub-phleo constructs allowing expression of the Hp/Hb receptor with native UTR sequences were generated in an identical fashion using the appropriate PCR primers. The Hp/Hb receptor alleles with native UTR sequences were transfected in *T. brucei* 427-060^R^ cells for analysis of mRNA stability.

### TLF Purification, Labelling, and Lysis Assays

Purified TLF was prepared as described in (53) and used in lysis assays. Briefly, 1 x 10^7^ cells were incubated at 37°C in a 300 μl volume with varying amounts of TLF. After 2 hrs incubation the percentage lysed cells were determined by hemocytometer counting. Each experiment was performed in triplicate. Alexa 488-Fluor labelled TLF, for FAC sorting, and Alexa 488-Fluor labelled Hp for binding studies were generated according to the manufacturer’s instructions (Invitrogen). Binding studies were performed as previously described (54).

### RT-PCR analysis

Total RNA was isolated with Tripure Isolation Reagent. cDNA was generated from 0.5 - 2 μg Turbo^TM^ DNase (ThermoFisher) treated total RNA with Superscript^TM^ III (ThermoFisher) according to the manufacturer’s instructions with oligo dT primers. Equal amounts of cDNA were used with Ready Go Taq Polymerase (Promega). To ensure specific DNA was amplified, a minus RT control reaction was used.

### Quantitative RT-PCR analysis

Total RNA was isolated and Turbo^TM^ DNase treated as described above. Quantification of Superscript^TM^ III generated cDNA was performed using an iCycler with an iQ5 multicolor real-time PCR detection system (Bio-Rad). Triplicate cDNA’s were analyzed and normalized to Asf I cDNA. qPCR oligonucleotide primers combos were designed using Integrated DNA Technologies software. cDNA reactions were diluted 10-fold and 5 µl was analyzed. A 15 µl reaction mixture contained 4.5 pmol sense and antisense primer, 7.5 µl 2X iQ SYBR green super mix (Bio-Rad Laboratories). Standard curves were prepared for each gene using 5-fold dilutions of a known quantity (100 ng/µl) of gDNA. The quantities were calculated using iQ5 optical detection system software.

### mRNA degradation experiments

Total RNA was obtained from *T. brucei* samples of 5×10^6^ cells taken at different times after the addition of 100μg/ml of Actinomycin D. Total RNA was extracted from each sample and qRT-PCR was performed using the method described above. mRNA half-lives were estimated after normalizing to 18S rRNA values and plotted on semi-logarithmic scales. Half-life of the mRNA is calculated as the amount of time required for a transcript to decrease 50% of initial amount. Half-life values were compared using Student’s t-test and a P value <0.001 was considered statistically significant.

### Nuclear run on

Nuclear run on was performed using 1×10^9^ cells as previously described (17,29). Briefly, parasites were washed twice at room temperature with 150 mM sucrose, 3-mM MgCl_2_, 20 mM KCl, 20 mM Hepes, 10-μg/ml leupeptin and 1mM dithiothreitol (buffer A) and resuspended in 800 μl of the same buffer. The parasite suspension was chilled on ice for 5 min, and palmitoyl-L-α-lysophosphatidylcholine (Sigma) dissolved in water at 10 mg/ml was added to obtain a final concentration of 250 μg/ml. After 1 min, the parasites were centrifuged, washed once with transcription buffer and incubated for 10 min on ice and 5 min at 30° C in the presence or absence of 500 ug/ml alfa-amanitin. Parasites were then resuspended in 200 μl of transcription labelling buffer, containing 2-mM adenosine triphosphate (ATP), 1-mM cytosine triphosphate (CTP), 1-mM guanosine triphosphate (GTP), 0.6-mg/ml creatine kinase (Roche Molecular Biochemicals), 25-mM creatine phosphate (Roche Molecular Biochemicals) and 0.75 mCi of [α-^32^P] uridine triphosphate (UTP) (6000 Ci/mmol). After 20 min incubation at 30° C, the parasites were centrifuged again; the pellets were lysed with 1 ml TriPure Isolation Reagent. The labeled RNA was recovered as described above. The labeled RNA was then hybridized in 5x saline/sodium phosphate/ethylenediaminetetraacetic acid (SSPE), 5 x Denhardt’s solution, 0.05 mg/ml yeast tRNA, 0.05 mg/ml Salmon Sperm DNA and 0.1% sodium dodecyl sulfate (SDS) to dot blots containing 2 μg spots of the indicated DNA. For preparation of the dot blots, PCR fragments containing probes for the targeted genes and regions was denatured in 0.2-N NaOH for 5 min at RT, neutralized by adding ammonium acetate to 1 N, and loaded onto nitrocellulose membranes using a minifold filtration apparatus. The 18S probe provide a control for RNAP I transcription and the 5S for RNAP III transcription. Tubulin probe provide control for RNAP II transcription. RNAs were incubated with membranes containing probes for 72 h at 65° C and then subjected to washing with 0.1 x SSC, 0.1% SDS three times at 65° C and then analyzed by phosphorimager.

### FAIRE analysis

Formaldehyde assisted regulatory elements (FAIRE) analysis was performed as previously described (16,50). Briefly, 1×10^7^ *T. brucei* cells were fixed in 1% formaldehyde for 5 min in HMI-9 media and reaction was terminated by adding 2.5M glycine to a final concentration of 125mM. Fixed cells were then lysed by adding a cell lysis solution containing 10mM EDTA, 50mMTris-HCL, 1% SDS and protease inhibitors. DNA was sonicated with a sonic amplitude sonicator for 10min (30-s on/off cycles) to obtain chromatin fragments of average 500bp in length. Debris were removed by centrifugation. No cross-linked sample was obtained for each replicate as total DNA control. DNA from both cross-linked and non-cross-linked samples were extracted by two consecutive phenol-chloroform extraction and DNA was ethanol precipitated in the presence of 20µg/ml of glycogen after RNA and proteinase K treatment. Each experiment was performed in triplicate and quantification of the FAIRE and total DNA samples were performed by real-time PCR and normalized to the analysis of 427.6.0740.

### ChIP analysis for Histone and Pol II

Chromatin immuno-precipitation (ChIP) was performed as previously described (16,50). Briefly, DNA was cross-linked to protein using formaldehyde as in FAIRE analysis. Sonicated DNA extract was pre-cleared using protein A agarose beads and incubated with or without relevant antibodies. Chromatin from 7×10^7^ cells equivalent was used in each immuno-precipitation reaction. Lysate was incubated with commercially prepared anti-histone H3 (AbCam) at a concentration of 2μg per IP reaction overnight at 4^0^C. *T. brucei* RNA Pol II antibody (a kind gift from Miguel Navarro) was used at 10μg per IP reaction. Protein-DNA complexes were incubated with protein agarose A beads for 2hrs and washed 3 times using wash buffers containing 0.1% SDS, 1% TritonX-100, 2mM EDTA, 20mM Tris, 500-150mM NaCL and protease inhibitors. DNA was eluted from beads using elution buffer containing 0.1% SDS and 0.1M NaHCO_3_. Cross-linking was then reversed by adding NaCl to a final concentration of 325 mM and incubated at 65°C overnight. DNA was then extracted using phenol-chloroform after RNAse and proteinase K treatments. Each ChIP experiment was performed in triplicate and analyzed by quantitative RT-PCR.

### Anti J dot-blot and immuno-precipitation

Anti J immunoblots were used to determine the global genomic levels of base J as previously described (55). Briefly, serially diluted genomic DNA was blotted onto nitrocellulose followed by incubation with anti-J antisera. Bound antibodies were detected with HRP conjugated goat anti-rabbit antibodies and visualized with ECL. DNA loading was determined by hybridization using a ^32^P-labeled tubulin probe. For the genome wide localization of base J, genomic DNA was sonicated and anti J immunoprecipitation performed as previously described (26). Each J-IP experiment was performed in triplicate and immunoprecipitated J containing DNA analyzed by quantitative PCR. Input DNA was used as a positive control for PCR.

### Strand-specific RNA-seq library construction and RNA-seq analysis

For mRNA-seq, total RNA was isolated from *T. brucei* cultures using TriPure. Three mRNA-seq libraries were constructed for each of the cell lines (*T. b. brucei* 427-221^S^, *T. b. brucei* 427-060^R^, and *T. b. brucei* 427-060^RS^) using Illumina TruSeq Stranded RNA LT Kit following the manufacturer’s instructions with limited modifications. The starting quantity of total RNA was adjusted to 1.3 µg, and all volumes were reduced to a third of the described quantity. High throughput sequencing was performed on an Illumina NovaSeq 6000 instrument. Raw reads from mRNA-seq were trimmed using Trim Galore! and locally aligned to the *T. brucei* Lister v427 genome assembly as previously described (37). Differential gene expression was conducted using DESeq2 (37) by comparing 427-221^S^ to 427-060^R^ and comparing 427-060^R^ to 427-060^RS^ in triplicate.

### DNA oligonucleotides

DNA oligos used in these studies are listed in Table S2.

### Data Availability

RNA-seq raw files and processed files have been deposited to the NCBI Gene Expression Omnibus (GEO) with accession number GSE273238.

## ACKNOWLEDGEMENTS

R.K. performed most of the experiments and analyzed data; L.C. executed the chromatin analyses; H.Y and R.J.S. carried out the RNA-seq analyses; R.S. and S.L.H. conceptualized and supervised the project and R.S. wrote the manuscript.

We are grateful for Erin Campbell for preparing the mRNA-seq libraries. We would like to thank Eric DeJesus for help with the TLF binding assays. We also like to thank Miguel Navarro for antibody against *T. brucei* Pol II.

This work was supported by the National Institutes of Health (grant number R01AI109108) to RS and RJS and (grant number AI39033) to SH. The funders had no role in study design, data collection, and interpretation, or the decision to submit the work for publication. The funders have not endorsed the work described here. The views reflected in the paper are solely those of the collaborating authors.

## SUPPLEMENTARY DATA

Supporting Information Table S1. Single nucleotide polymorphisms (SNPs) in the 427 T. brucei genome for (A) Tb427.6.440; (B) Tb427.6.360; (C) Tb427.6.650; and (D) Tb427.6.740. The nucleotide position and the SNP identified in the 427 *T. brucei* genome is provided along with confirmation in genomic DNA PCR in the parental 427-221^S^ cells. Results of direct sequencing of RT-PCR mRNA analysis are presented for the 427-VO2^SR^ and 427-060^RS^ cells are indicated. N.A, not analyzed

Supporting Information Table S2. Oligos used in these studies.

