## Supplemental Figures for "Mono-allelic epigenetic regulation of bi-directional silencing of RNA Polymerase II polycistronic transcription initiation in *Trypanosoma brucei*"

### Fig S1

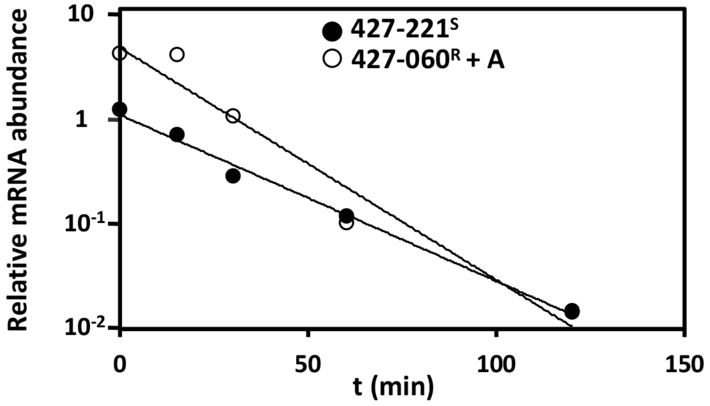


**Supplemental Figure 1. Half-life of HpHbR mRNA is not altered in TLF resistant T. b. brucei 427-060$^{R}$ cells.** Actinomycin-D treatment of *T. b. brucei* 427-221$^{S}$ (black circle) and *T. b. brucei* 427-060$^{R}$ cells, ectopically expressing Tb427.6.440 allelic copy A (white circle) from the tubulin locus. HpHbR mRNA levels are normalizes versus 18S rRNA.

### Fig S2

**A**

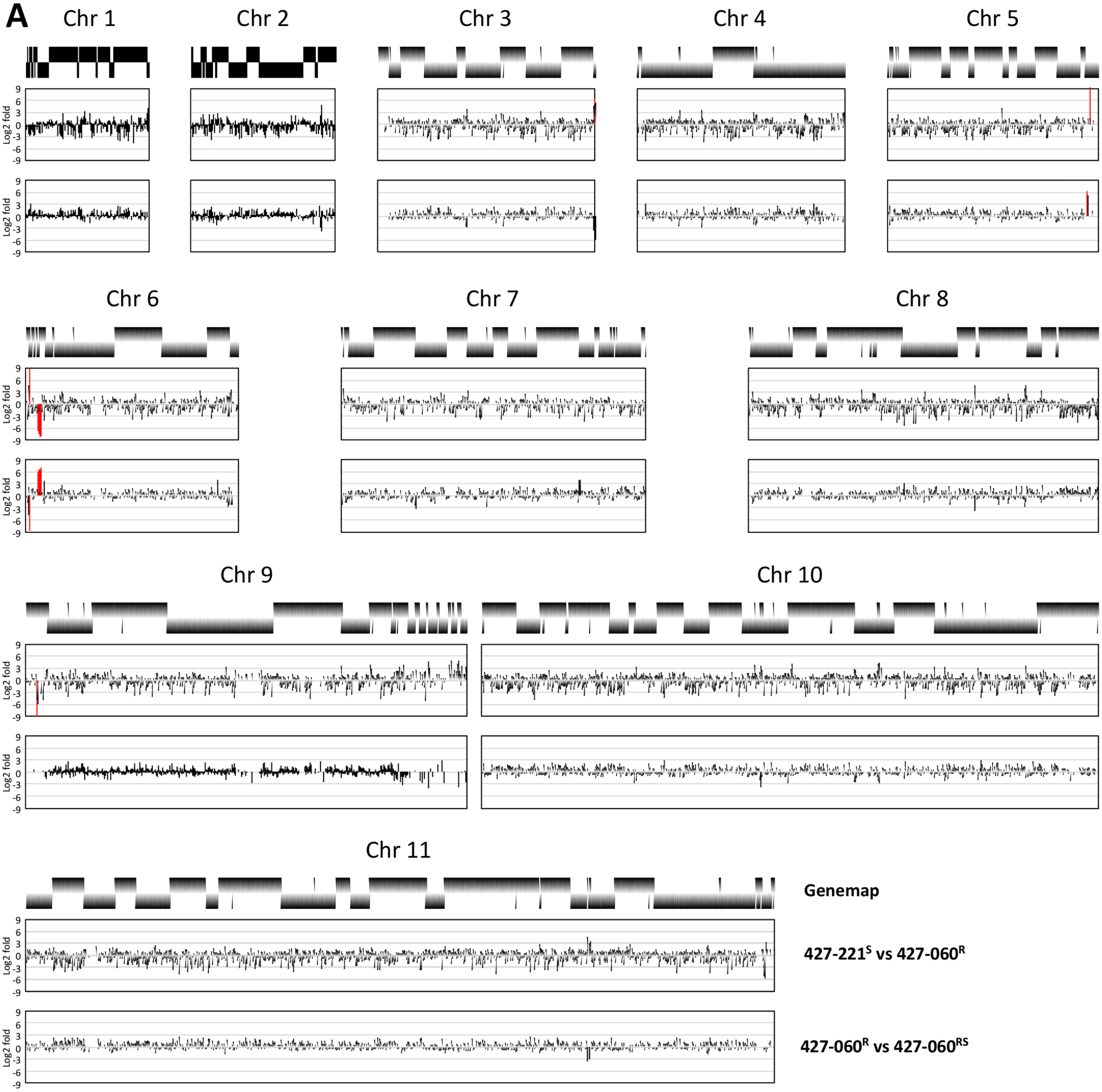


**B**

| Gene ID | 427-221$^{S}$ vs 427-060$^{R}$ Log2Fold Change | 427-060$^{R}$ vs 427-060$^{RS}$ Log2Fold Change | Annotation |
|---|---|---|---|
| Tb427.03.5880 | 6.41 | -3.62 | variant surface glycoprotein, point mutation |
| Tb427.05.4750 | -0.32 | 6.31 | variant surface glycoprotein, degenerate |
| Tb427.05.4980 | 11.63 | -0.02 | variant surface glycoprotein, frameshift (VSG 060) |
| Tb427.06.180 | 8.83 | -8.64 | receptor-type adenylate cyclase GRESAG 4, putative |
| **Tb427.06.360** | **-6.68** | **6.06** | **UDP-Gal or UDP- GlcNAc-dependent glycosyltransferase, degenerate** |
| **Tb427.06.400** | **-7.55** | **6.49** | **peptidase M20/M25/M40, putative** |
| **Tb427.06.410** | **-8.16** | **6.63** | **hypothetical protein%2C conserved** |
| **Tb427.06.420** | **-6.92** | **6.26** | **hypothetical protein** |
| **Tb427.06.430** | **-7.31** | **6.48** | **receptor-type adenylate cyclase GRESAG 4%2C putative** |
| **Tb427.06.440** | **-8.00** | **7.13** | **haptoglobin-hemoglobin receptor** |
| Tb427tmp.142.0130 | -9.24 | N.D. | variant surface glycoprotein (VSG%, pseudogene), putative |

**Supplemental Figure 2. Genome-wide view of gene expression changes associated with silencing of HpHbR expression in TLF resistant *T. brucei*.**

**(A)** Changes in transcript levels based on RNA-seq reads during *in vitro* TLF selection and generation of *T. b. brucei* 427-060$^{R}$ and 427-060$^{RS}$ cell lines are shown for all chromosomes. Gene map of for each chromosome is shown above. mRNA coding genes on the top strand are indicated by black lines in the top half of the panel, bottom strand by a line in the bottom half. Genes on the top strand are transcribed from left to right and those on the bottom strand are transcribed from right to left. Plots representing the log2 fold changes that occur during conversion of *T. b. brucei* 427-221$^{S}$ to *the* resistant 427-060$^{R}$ and during conversion of TLF resistant *T. b. brucei* 427-060$^{R}$ to sensitive 427-060$^{RS}$ are presented as indicated. Genes with altered expression using a cut-off value of a 6-fold log2 are highlighted in red. **(B)** List of all genes with altered expression levels greater than 6-fold log2.

#### Fig S3

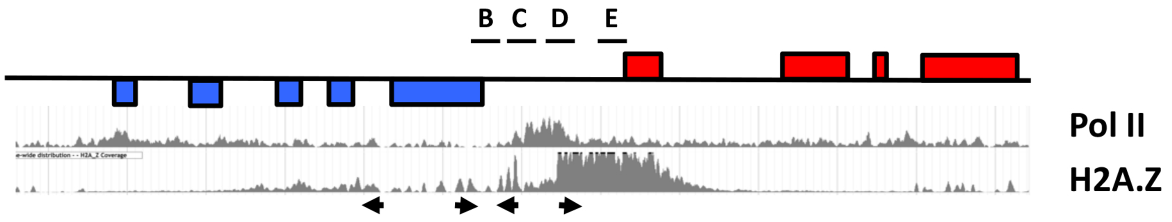


**Supplemental Figure 3. ChIP-seq data of the RNA Pol II subunit RBP9 and H2A.Z**

Data from [22] is shown across the divergent SSR representing the transcription start region of the divergent PTUs on chr 6 as shown in Figure 4. Arrowheads below represent the transcription start sites for each PTU mapped by small 5'-triphosphate-RNA-seq. The apparent single peak of Pol II is presumably due to the small size of this dSSR limiting the resolving power of the Pol II ChIP.
