## Supplemental Tables for "Mono-allelic epigenetic regulation of bi-directional silencing of RNA Polymerase II polycistronic transcription initiation in *Trypanosoma brucei*"

**Table S1A.** *SNP analysis Tb427.6.440 (pos. 1 - 1212)*

| Genomic position | ORF position | Tb-427 genome | 427-221$^{S}$ | 427-VO2$^{SR}$ (allele A) | 427-060$^{RS}$ (allele B) |
|---|---|---|---|---|---|
| 206507 | 261 | G / C | G / C | G | C |
| 206815 | 569 | C / T | C / T | C | T |
| 206819 | 573 | A / G | A / G | A | G |
| 207124 | 878 | C / T | C / T | C | T |
| 207172 | 926 | G / A | G / A | G | A |
| 207216 | 970 | C / T | C / T | C | T |
| 207352 | 1106 | A / G | A / G | A | G |
| 207355 | 1109 | G / C | G / C | G | C |

**Table S1B.** *SNP analysis Tb427.6.360 (pos. 1 - 1212)*

| Genomic position | ORF position | Tb-427 genome | 427-221$^{S}$ | 427-VO2$^{SR}$ (allele A) | 427-060$^{RS}$ (allele B) |
|---|---|---|---|---|---|
| 176626 | 273-275 | TGT | TGT | TGT | TGT |
| 176755 | 142 | C / T | C / T | T | C |
| 176799 | 98 | C / A | C / A | C | A |

**Table S1C.** *SNP analysis Tb427.6.650 (pos. 1 - 2400)*

| Genomic position | ORF position | Tb-427 genome | 427-221$^{S}$ | 427-VO2$^{SR}$ | 427-060$^{RS}$ |
|---|---|---|---|---|---|
| 276650 | 1512 | A / G | A / G | A / G | A / G |
| 276703 | 1459 | A / G | A / G | A / G | A / G |
| 276982 | 1180 | T / C | N.A. | N.A. | N.A. |
| 277010 | 1152 | T / C | N.A. | N.A. | N.A. |
| 277022 | 1140 | A / T | N.A. | N.A. | N.A. |
| 277132 | 1030 | T / C | N.A. | N.A. | N.A. |
| 277147 | 1015 | C / T | C / T | C / T | C / T |
| 277199 | 963 | A / C | A / C | A / C | A / C |
| 277308 | 854 | A / G | A / G | A / G | A / G |

**Table S1D.** *SNP analysis Tb427.6.740 (pos. 1 - 2400)*

| Genomic position | ORF position | Tb-427 genome | 427-221$^{S}$ | 427-VO2$^{SR}$ | 427-060$^{RS}$ |
|---|---|---|---|---|---|
| 305014 | 2185 | G / T | G / T | G / T | G / T |
| 306206 | 993 | T / C | T / C | T / C | T / C |
| 306755 | 444 | A / G | A / G | A / G | A / G |

**Table S2.** *Oligonucleotides used*

| Name | Orientation | Sequence |
|---|---|---|
| Run on fragment 1 | sense | 5’-TTACATAATCGAGCATCCCGC-3’ |
| Run on fragment 1 | antisense | 5’-ACTGATGGTATCGTCAGGTT-3’ |
| Run on fragment 2 | sense | 5’-TCACGAAGCCACTACATGCG-3’ |
| Run on fragment 2 | antisense | 5’-ATGCTACTTCCCCTCTACCTA-3’ |
| Run on fragment 3 | sense | 5’-AGGTTCATCACATCTCTTCGC-3’ |
| Run on fragment 3 | antisense | 5’-AACAGAGGAAAACACGATGCG-3’ |
| Run on fragment 4 | sense | 5’-CTAGCATGTAATTCACGCAT-3’ |
| Run on fragment 4 | antisense | 5’-GCATTGATCAGTCCCATCTC-3’ |
| Run on fragment 5 | sense | 5’-CGGTTGGTTTTTCTCTTCAG-3’ |
| Run on fragment 5 | antisense | 5’-CATGAGTGAAAGACACACGA-3’ |
| Run on fragment 6 | sense | 5’-TTTCTTTGTGTTACAAGGGG-3’ |
| Run on fragment 6 | antisense | 5’-CTGAAACTATCATTATTACC-3’ |
| Run on fragment 7 | sense | 5’-ACCCAGGCCATGCTTCCAGA-3’ |
| Run on fragment 7 | antisense | 5’-GCACCCCAATCAACCTGCAT-3’ |
| Run on fragment 8 | sense | 5’-ATGCAGGTTGATTGGGGTGCA-3’ |
| Run on fragment 8 | antisense | 5’-TCACTGCTTATGTATCTTTGG-3’ |
| Run on fragment 9 | sense | 5’-GGTCCATTCCTTTTATTGCAATG-3’ |
| Run on fragment 9 | antisense | 5’-CACCGATGTAACTGAGCACAG-3’ |
| Run on fragment 10 | sense | 5’-ATGACACAACCAGATATATTC-3’ |
| Run on fragment 10 | antisense | 5’-CTGAAGGGTCTTAAGGGAAG-3’ |
| Run on fragment 11 | sense | 5’-CTACATTACCAACAAATCTC-3’ |
| Run on fragment 11 | antisense | 5’-TCAGCATCTGTAACGAACTAC-3’ |
| Run on fragment 12 | sense | 5’-CGAGGGTCCGAAAGTTTTGTA-3’ |
| Run on fragment 12 | antisense | 5’-GGAGCTGCTTTGGAGCCTTGT-3’ |
| Run on fragment 13 | sense | 5’-GAATGTTGCTGCTACGGTATG-3’ |
| Run on fragment 13 | antisense | 5’-ATCGTCTAGACAAAGCTGCGACTGCACCCC-3’ |
| Run on fragment 14 | sense | 5’-GATCCCTGCAGGATGGAGAAACCGTCTTGCAG-3’ |
| Run on fragment 14 | antisense | 5’-GATCGGCGCGCCCTACACCACCACCTGGAGCA-3’ |
| Run on fragment 15 | sense | 5’-CTACGCCGTGTTGGGTGCAAG-3’ |
| Run on fragment 15 | antisense | 5’-GTCGCCACGGGGTCTACATGC-3’ |
| Run on fragment 16 | sense | 5’-AGTCAGAGACGGGGGTCCCT-3’ |
| Run on fragment 16 | antisense | 5’-ATGTTGGCCACACGTCAGATA-3’ |
| Run on fragment 17 | sense | 5’-CGGATCCAAAACACCAAAACA-3’ |
| Run on fragment 17 | antisense | 5’-ATGAACCAGAAGCGATGCGAA-3’ |
| Run on fragment 18 | sense | 5’-ATGCGCGAAATCGTCTGCGTTC-3’ |
| Run on fragment 18 | antisense | 5’-CTAGTATTGCTCCTCCTCGTC-3’ |
| Run on fragment 19 | sense | 5’-GGGTACGACCATACTTGGCCG-3’ |

| | | |
|---|---|---|
| Run on fragment 19 | antisense | 5’-GAGTACAACACCCCGGGTTCC-3’ |
| Run on fragment 20 | sense | 5’-GTCATATGCTTGTTTCAAG-3’ |
| Run on fragment 20 | antisense | 5’-GACTTTTGCTTCCTCTATTG-3’ |
| qPCR fragment A | sense | 5’-TAGGCAGTGAGGCGTTCGTTAGTT-3’ |
| qPCR fragment A | antisense | 5’-TTGGTAAGGTGTACCCAACTGCCT-3’ |
| qPCR fragment B | sense | 5’-ATTTGGATGCAGTGCGACCACACGAA-3’ |
| qPCR fragment B | antisense | 5’-CTGCTGCAAAGGATTGTCGCTGTA’3’ |
| qPCR fragment C | sense | 5’-CAGAACGCTTGTTTGCCCTCCAAT-3’ |
| qPCR fragment C | antisense | 5’-AAACCGTTCACTCCGCCCTTAGTT-3’ |
| qPCR fragment D | sense | 5’-TGTCTCCTTGTGTTCACTTGTGGG-3’ |
| qPCR fragment D | antisense | 5’-AACTTCCACGCACCTTTGCTACCT-3’ |
| qPCR fragment E | sense | 5’-CTCGGCTATTGTATTTGGCCGCAA-3’ |
| qPCR fragment E | antisense | 5’-TTGCACCTGCACTTTCCTCCTCTA-3’ |
| qPCR fragment F | sense | 5’-TTTGTGGTGTTACGGCACCTGTTG-3’ |
| qPCR fragment F | antisense | 5’-AGTGTCGTTCAGTGGTCCGGTAAA-3’ |
| qPCR Tb927.6.740 | sense | 5’-TGTATTGTGGCGGACATGACGACT-3’ |
| qPCR Tb927.6.740 | antisense | 5’-TCTCATTGTGGATGAGGTGCACGA-3’ |
| qPCR SL promoter | sense | 5’-CTTTGTTTCCCATAAGTCTAC-3’ |
| qPCR SL promoter | antisense | 5’-AGACACTTGCCATATTTTACT-3’ |
| Splice leader RNA | sense | 5’-CGCTATTATTAGAACAGTTTCTGTAC-3’ |
| qPCR enolase | antisense | 5’-TCTCTGTCGTCACTTCAACCTCCA-3’ |
| qPCR HpHbR | antisense | 5’-CGCCTTCTCAACTTCGTCTTTGGT-3’ |
| RT Tb927.6.350 | sense | 5’-ATGACGATTCAAGCCGTATCG-3’ |
| HpHbR EcoR I | sense | 5’-GATCGAATTCATGGAGAAACCGTCTTGCAG-3’ |
| HpHbR EcoR I | antisense | 5’-GATCGAATTCCTACACCACCACCTGGAGCA-3’ |
| HpHbR seq | sense | 5’-TTTGGGTTCTGCTTAATGCC-3’ |
| HpHbR seq | antisense | 5’-TTAGACAATTTAAACTTGTTCAGC-3’ |

Restriction sites used for cloning are underlined.
